# Strong homeostatic TCR signals induce formation of self-tolerant virtual memory CD8 T cells

**DOI:** 10.1101/202218

**Authors:** Ales Drobek, Alena Moudra, Daniel Mueller, Martina Huranova, Veronika Horkova, Michaela Pribikova, Robert Ivanek, Susanne Oberle, Dietmar Zehn, Kathy D. McCoy, Peter Draber, Ondrej Stepanek

**Affiliations:** Laboratory of Adaptive Immunity, Institute of Molecular Genetics, Czech Academy of Sciences, 14220 Prague, Czech Republic; Department of Biomedicine, University Hospital and University of Basel, 4031 Basel, Switzerland; Swiss Institute of Bioinformatics, Basel, Switzerland; Swiss Vaccine Research Institute, 1066 Epalinges, Switzerland; Division of Animal Physiology and Immunology, School of Life Sciences Weihenstephan, Technical University of Munich, 85354 Freising, Germany; Department of Clinical Research (DKF), University of Bern Inselspital, 3012 Bern, Switzerlands

## Abstract

Virtual memory T cells are foreign antigen-inexperienced T cells that have acquired memory-like phenotype and constitute for 10-20% of all peripheral CD8^+^ T cells in mice. Their origin, biological roles, and relationship to naïve and foreign antigen-experienced memory T cells are incompletely understood. By analyzing TCR repertoires and using retrogenic monoclonal T-cell populations, we show that virtual memory T cells originate exclusively from strongly self-reactive T cells. Moreover, we show that the stoichiometry of the CD8 interaction with Lck regulates the size of the virtual memory T-cell compartment via modulating the self-reactivity of individual T-cell clones. We propose a so far unappreciated peripheral T-cell fate decision checkpoint that eventually leads to the differentiation of highly self-reactive T cells into virtual memory T cells. This underlines the importance of the variable level of self-reactivity in polyclonal T cells for the generation of functional T-cell diversity. Although virtual memory T cells descend from the highly self-reactive clones and acquire a partial memory program, they do not show higher capacity to induce autoimmune diabetes than naïve T cells. Thus, virtual memory T cells are not generally more responsive than naïve T cells, because their activity highly depends on the immunological context.

**Summary:** We conclude that virtual memory T cells are formed from self-reactive CD8^+^ T cells in a process regulated by CD8-Lck stoichiometry. Despite their self-reactivity and partial memory differentiation program, virtual memory T cells did not show a strong autoimmune potential.

## Introduction

Immunological memory is one of the hallmarks of adaptive immunity. During infection, pathogen-specific naïve T cells differentiate into short lived effector and memory T cells. The latter facilitate long-standing protection against a secondary infection of the same pathogen. CD8^+^ CD44^+^ CD62L^+^ central memory (CM) T cells have the capability of expansion, self-renewal, and generation of cytotoxic effector T cells upon repeated encountering of their cognate antigen [1].

Interestingly, some T cells with an apparent memory phenotype are specific to antigens which the host organism has not been exposed to [2, 3]. There are two possible explanations of their origin: (i) they could be cross-reactive T cells that have encountered another foreign cognate antigen previously [3] or (ii) they are generated via homeostatic mechanisms independently of the exposure to any foreign antigens [2]. The strong evidence for the role of homeostatic mechanisms in generation of CD8^+^ memory-like T cells was provided by the detection of these cells in germ-free mice. Since these T cells had limited prior exposure to foreign antigens, they were called virtual memory (VM) T cells [2].

Generation and/or maintenance of VM T cells depends on transcription factors Eomes and IRF4, type I interferon signaling, IL-15 and/or IL-4 signaling, and CD8α^+^ dendritic cells [4–8]. It has been shown that VM T cells express slightly higher levels of CD122 (IL-2Rβ) and lower levels of CD49d (integrin α4; a subunit of VLA-4) than true (i.e., foreign antigen-experienced) CM T cells [4]. Based on these markers, VM T cells constitute for a majority of memory-phenotype CD8^+^ T cells in unimmunized mice and around 10-20% of total CD8^+^ T cells in lymphoid organs. Moreover, it has been proposed that memory-phenotype T cells, that accumulate in aged mice, are VM T cells [9]. However, with the notable exception of the very initial study that identified CD44^+^ CD62L^+^ CM T cells in germ-free mice [2], all other published experiments used specific pathogen free (SPF) mice, that have significant exposure to microbial antigens. There are three major hypothesis of how virtual memory T cells might be formed in a homeostatic manner: (i) the differentiation into VM T cells occurs purely on a stochastic basis, (ii) lymphopenic environment in newborns induces differentiation of the first wave of thymic emigrants into VM T cells [8], (ii) virtual memory T cells are formed on a purely stochastic basis, or (iii) the VM T cells are generated from relatively highly self-reactive T cells that receive strong homeostatic TCR signals at the periphery [5]. However, none of these hypotheses has been addressed in detail.

Although VM T cells form a large CD8^+^ T cell population, their biological role is still unknown. VM T cells share some functional properties with true CM cells, including rapid production of IFNγ upon stimulation with a cognate antigen or cytokines [2, 10]. On a per cell basis, ovalbumin-specific VM T cells provide a protection to ovalbumin-expressing Listeria monocytogenes (Lm), comparable to true CM T cells and surpass naïve T cells with the same specificity [10]. In addition, it has been proposed that VM T cells are capable of by-stander protection against infection, i.e. independently of their cognate antigen exposure [5, 11]. Altogether these data pointed towards the superior role of VM cells in protective immunity to infection. In a marked contrast to the above mentioned findings, Decman et al. showed that CD44^+^ CD8^+^ T-cell receptor (TCR) transgenic T cells isolated from unprimed mice (i.e. putative VM T cells) expand less than CD44^-^ CD8^+^ T cells expressing the same TCR upon antigenic stimulation in vivo [12]. Likewise, VM T cells from aged mice were shown to be hyporesponsive to their cognate antigens in comparison to their naïve counterparts, mostly because of their susceptibility to apoptosis [12, 13].

As VM T cells develop independently of infection, the understanding of mechanisms that guide their development is critical in order to elucidate their biological roles. One hint is the observation that levels of CD5 (a marker of self-reactivity) on naïve T cells correlate with their ability to differentiate into VM T cells. [5]. In this study, we demonstrate that virtual memory T cells originate exclusively from relatively highly self-reactive T-cell clones and acquire only a partial memory gene-expression
program. Moreover, the interaction between CD8 and Lck (and possibly the overall intrinsic sensitivity of the TCR signaling machinery) determines the size of the virtual memory compartment. These data highlight the virtual memory T-cell formation as a T-cell fate decision checkpoint, when the intensity of TCR signals induced by self-antigens plays a central role in the decision-making process. Although virtual memory T cells show augmented responses to their foreign cognate antigen in some experimental setups in comparison to naïve T cells, the capability to induce autoimmune diabetes is comparable in naïve and virtual memory T cells.

## Results

### Strong homeostatic TCR signaling induces virtual memory T cells

In this work, we aimed to understand why some mature CD8^+^ T cells differentiate into VM T cells and some maintain their naïve phenotype. The peripheral T-cell pool of polyclonal mice consists of clones with different level of self-reactivity. To test the hypothesis that the level of self-reactivity plays a role in the formation of virtual memory T cells, we took advantage of previous observations that coupling frequency (or stoichiometry) of CD8 coreceptor to Lck kinase is a limiting factor for the TCR signaling in thymocytes [14, 15]. We used T cells from CD8.4 knock-in mouse strain, that express a chimeric CD8.4 coreceptor, consisting of the extracellular portion of CD8 fused to the intracellular part of CD4 [15]. In comparison to WT CD8α, the chimeric CD8.4 coreceptor is more strongly binding Lck, a kinase initiating the TCR signal transduction. As a consequence, higher fraction of CD8.4 coreceptors molecules than CD8 coreceptors are loaded with Lck [14, 15]. Because the self-antigen triggered TCR signaling is stronger in CD8.4 T cells than CD8WT T cells [14, 16, 17], we use the CD8.4 T cells as a model to address the role of self-reactivity in virtual memory T-cell formation.

First, we compared monoclonal F5 Rag^−/−^ T cells (henceforth CD8WT F5) and CD8.4 knock-in F5 Rag^−/−^ T cells (henceforth CD8.4 F5) from unimmunized animals. The F5 TCR is specific for influenza NP68 and has been shown to have a very low level of self-reactivity [18, 19]. We observed that CD8.4 F5 T cells had lower levels of CD8 and TCR and elevated levels of CD5 and IL-7R in comparison to CD8WT F5 T cells (Fig. S1A). Because the downregulation of CD8 and expression of CD5 and IL-7R correlate with the intensity of homeostatic TCR signaling [17], we could conclude that CD8.4 indeed enhances homeostatic TCR signaling. However, we did not detect upregulation of memory markers, CD44, CD122, and LFA-1 on CD8.4 F5 T cells (Fig. 1A, S1A). CD8.4 F5 T cells showed slightly stronger antigenic responses, measured as CD25 and CD69 upregulation, than CD8WT F5 T cells in vitro upon activation with the cognate antigen, NP68, or a lower affinity antigen, NP372E [20] (Fig. 1B). Accordingly, CD8.4 F5 T cells expanded more than CD8WT F5 T cells after the immunization with NP68 peptide (Fig. 1C). Infection with transgenic Listeria monocytogenes expressing NP68 (Lm-NP68) induced stronger expansion and formation of larger KLRG1^+^IL-7R^−^ effector and KLRG1^−^IL-7R^+^ memory-precursor subsets in CD8.4 F5 than in CD8WT F5 T cells (Fig. 1D, Fig. S1B). Collectively, these data showed that CD8-Lck coupling frequency sets the sensitivity of peripheral T cells to self-antigens during homeostasis and to foreign cognate antigens during infection. However, supraphysiological CD8-Lck coupling in CD8.4 F5 T cells, does not induce differentiation into memory-phenotype T cells in unimmunized mice.

**Figure 1.**
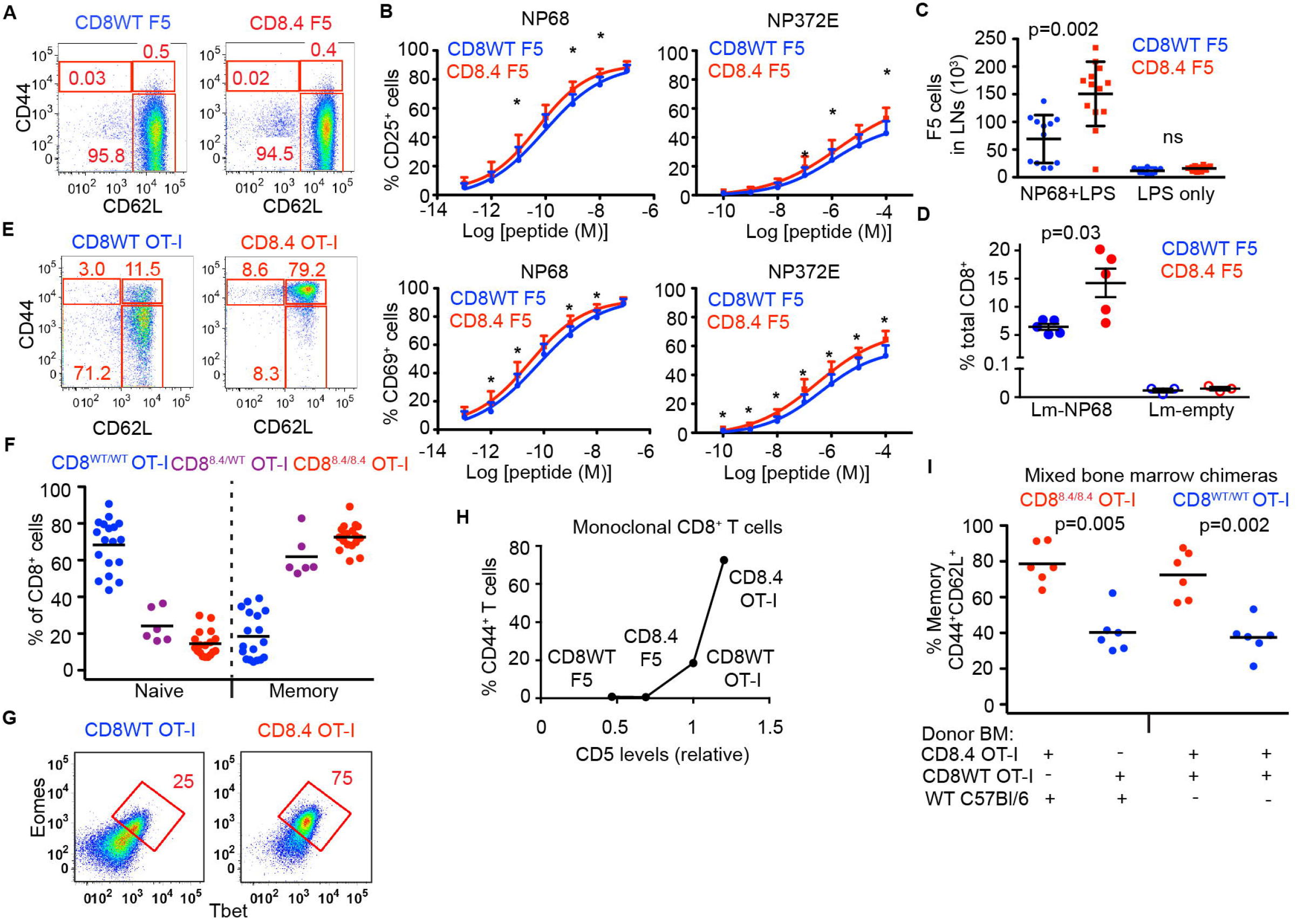
Supraphysiological CD8-Lck coupling induces differentiation into VM T cells in a clone-specific manner. (A) LN cells isolated from CD8WT F5 and CD8.4 F5 mice were analyzed by flow cytometry (gated as viable CD8^+^CD4^−^). A representative experiment out of 4 in total. (B) LN cells isolated from CD8WT F5 and CD8.4 F5 mice were stimulated with antigen-loaded DCs overnight and CD69 or CD25 expressionwas analyzed by flow cytometry. Mean + SEM. n=7 independent experiments. (C) 2×10^6^ CD8WT F5 or CD8.4 F5 LN T cells were adoptively transferred into Ly5.1 WT host 1 day prior to immunization with NP68 peptide and LPS or LPS only. 3 days after the immunization, donor Ly5.2^+^ Ly5.1^−^ CD8^+^ T cells from LN of the host mice were analyzed by flow cytometry and counted. Median ± interquartile range. n=8-13 mice from 7 independent experiments. (D) 1×10^5^ CD8WT F5 or CD8.4 F5 LN T cells were adoptively transferred into Ly5.1 WT hosts and immunized with WT Lm (empty) or Lm-NP68. 6 days after the immunization, the percentage of donor Ly5.2^+^ Ly5.1^−^ CD8^+^ T cells among all CD8^+^ T cells was determined. n=3-5 mice from 3 independent experiments. (E-F) LN cells isolated from CD8WT OT-I, CD8.4 OT-I, and heterozygous CD8^WT/8.4^ mice were analyzed by flow cytometry (gated as viable CD8^+^ CD4^−^). Percentages of naïve (CD44^−^ CD62L^+^) and memory (CD44^−^ CD62L^+^) CD8^+^ T cells were determined. n=6-18 mice from at least 5 independent experiments. (G) Expression of Eomes and Tbet in CD8^+^ LN T cells isolated from CD8WT OT-I and CD8.4 OT-I mice was determined by flow cytometry. A representative experiment out of 3 in total. (H) Relationship between relative CD5 levels (CD5 on CD8WT OT-I was arbitrarily set as 1) percentage of VM T cells (CD44^+^ CD62L^+^) using the data from indicated TCR transgenic strains. Mean value of n=5-8 mice from at least 5 independent experiments. (I) Irradiated Rag2^−/−^ host mice were transplanted with B-cell and T-cell depleted bone marrow from Ly5.1 C57Bl/6 together with CD8.4 OT-I or CD8WT OT-I bone marrow in 1:1 ratio (first two datasets) or, with bone marrow from CD8.4 OT-I and CD8WT OT-I mice in 1:1 ratio (last two datasets). 8 weeks later, LN cells were isolated and analyzed by flow cytometry. n=6 mice from 3 independent experiments. Statistical significance was determined using 2-tailed Mann-Whitney test (B-D, I). * p < 0.05

Whereas the level of self-reactivity of F5 T cells is very low, transgenic OT-I T cells, specific for chicken ovalbumin (OVA), exhibit a relatively high level of self-reactivity [18, 19]. We tested whether a combination of a relatively highly self-reactive OT-I TCR and supraphysiological CD8-Lck coupling is sufficient to induce VM T cells. For this reason, we compared monoclonal OT-I Rag^−/−^ T cells (henceforth CD8WT OT-I) and CD8.4 knock-in OT-I Rag^−/−^ T cells (henceforth CD8.4 OT-I) from unimmunized animals. As expected, CD8.4 OT-I T cells exhibited signs of stronger homeostatic TCR signaling than CD8WT OT-I T cells, including downregulation of TCR, CD8, and increased levels of CD5 and CD127 (Fig. S1C-D). Interestingly, the majority of the CD8.4 OT-I T cells exhibited CM phenotype including CD44^+^ CD62L^+^ double positivity, increased levels of CD122 and LFA-1, increased forward-scatter signal, and expression of transcription factors Tbet and Eomes (Fig. 1E-G, S1C), which was in a striking contrast with analogical experiments with CD8WT/CD8.4 F5 T cells (Fig. 1A, S1A). The CD8^WT/CD8.4^ heterozygous OT-I T cells showed an intermediate frequency of memory T cells (Fig. 1F). Because CD8.4 OT-I T cells exhibited features of memory T cells without encountering their foreign cognate antigen, we concluded that CD8.4 induced a differentiation of OT-I T cells into VM cells. When we compared monoclonal T cells from all four transgenic mouse strains tested, we identified a relationship between surface levels of CD5, a commonly used marker of self-reactivity, and the frequency of VM T-cell formation. However, the dependency was not linear, but showed a threshold behavior, indicating that only the most self-reactive T cells have the potential to develop into VM T cells (Fig. 1H, S1E).

To address whether CD8.4 induces VM T-cell formation in OT-I T cells in a T-cell intrinsic manner, we generated mixed bone marrow chimeric animals where both CD8.4 and CD8WT populations were present. We transplanted bone marrow cells from Ly5.1 WT mouse together with bone marrow cells from either Ly5.2 CD8WT OT-I or Ly5.2 CD8.4 OT-I mice into an irradiated Rag2^−/−^ recipient. CD8.4 OT-I T cells generated substantially more VM T cells than CD8WT OT-I T cells (Fig. 1I). In an alternative setup, we co-transferred mixed bone marrow cells from Ly5.1 CD8WT OT-I and Ly5.2 CD8.4 OT-I animals into an irradiated Rag2^−/−^ recipient to compare these two subsets in a single mouse. Again, CD8.4 OT-I T cells generated significantly more VM T cells than CD8WT OT-I T cells (Fig. 1I). These experiments showed that CD8.4 OT-I T-cells intrinsically trigger the memory differentiation program.

### CD8-Lck coupling frequency regulates the size of virtual memory compartment in polyclonal repertoire

In a next step, we addressed how the CD8-Lck stoichiometry regulates the size of virtual memory compartment in a polyclonal T-cell pool. Interestingly, CD8.4 polyclonal mice showed significantly higher frequency of VM CD8^+^ T cells than CD8WT control animals, although most CD8.4 T cells still showed a naïve phenotype (Fig. 2A-B). Thus, CD8.4 induced the VM T-cell formation only in a subset of polyclonal CD8^+^ T cells, implying that enhanced CD8-Lck coupling has clone-specific effects. The VM T cells from both CD8WT and CD8.4 mice rapidly produced IFNγ after stimulation with PMA/ionomycine, showing that CD8WT and CD8.4 VM T cells are indistinguishable in this functional trait, typical for memory T cells (Fig. S2A). Importantly, the analysis of mice in germ-free condition showed elevated frequency of CD44^+^ and T-bet/Eomes double-positive T cells in CD8.4 mice when compared to CD8WT (Fig. 2C-D), demonstrating that supraphysiological CD8-Lck coupling indeed promotes formation of VM T cells independently of the exposure to foreign antigens.

**Figure 2.**
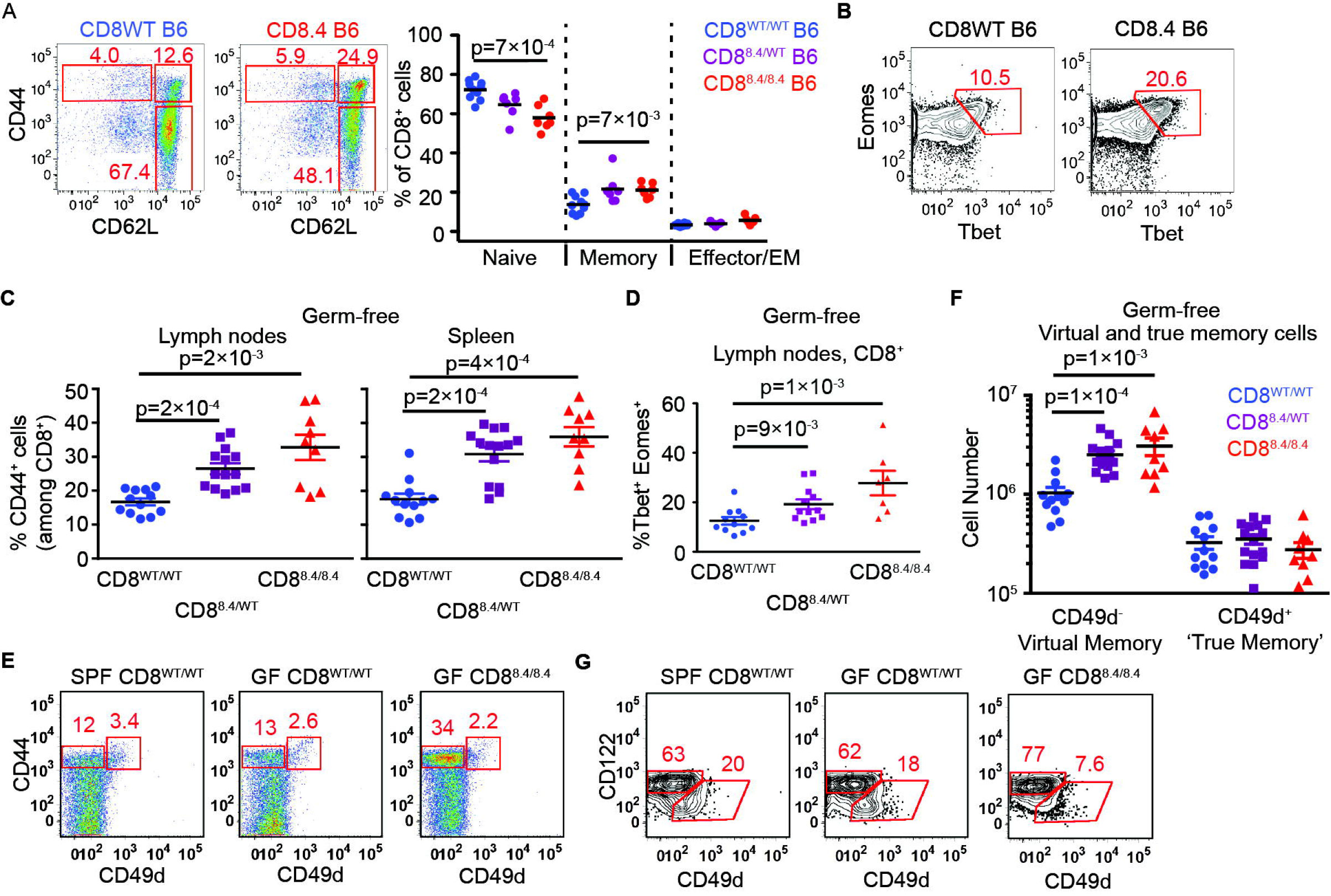
CD8-Lck coupling is a limiting factor for the size of the virtual memory T cell compartment. (A-B) Percentages of naïve (CD44^−^CD62L^+^), central memory (CD44^+^ CD62L^+^), and effector/effector memory (CD44^+^CD62L^−^) (A) and Eomes^+^ Tbet^+^ (B) CD8^+^ LN T cells isolated from CD8WT and CD8.4 mice were determined by flow cytometry. Representative experiments out of 7 (A) or 5 (B) in total. (C-F) LN cells and splenocytes were isolated from germ-free polyclonal CD8WT, CD8.4, and heterozygous CD8^WT/8.4^ mice and CD8^+^ T cells were analyzed by flow cytometry. (C) Percentage of CD44^+^ T cells among CD8^+^ LN cells and splenocytes. Mean + SEM. n=9-14 mice from 4 independent experiments. (D) Percentage of Tbet^+^ Eomes^+^ double-positive T cells. Mean ° SEM. n=7-12 mice from 3 independent experiments. (E) Percentage of CD44^+^ CD49d^−^ VM and CD44^+^ CD49d^+^ true memory T cells in the spleen. A representative experiment out of 4 in total. (F) Absolute numbers of CD8^+^ CD44^+^ CD49d^-^ VM and CD8^+^ CD44^+^ CD49d^+^ true memory T cells in LN and the spleen were quantified. n=9-14 mice from 4 independent experiments. (G) Percentage of CD122^HI^ CD49d^-^ VM and CD122^LOW^ CD49^+^ true CM cells among CD8^+^ CD44^+^ CD62L^+^ CM T cells isolated from LN. A representative experiment out of 3 in total. Statistical significance was determined using 2-tailed Mann-Whitney test.

Although the VM and true CM T cells are very similar in many aspects, VM T cells were previously reported to express lower levels of CD49d and slightly higher levels of CD122 than true CM T cells [2, 4, 5, 9]. However, CD49d as a marker discriminating VM and true CM T cells has not been validated using T cells from germ-free animals. For the first time, we could show that CD49d^−^ and CD49d^+^ T cells within the CD44^+^ compartment of SPF and germ-free mice occur at comparable frequencies (Fig. 2E). Only the CD49d^−^ CD44^+^, but not the CD49d^+^ CD44^+^, subset was expanded in the CD8.4 mouse, indicating that these subsets are not related (Fig. 2E). Accordingly, CD122^HI^ CD49d^−^ CD44^+^ T cells, but not CD122^LOW^ CD49d^+^ CD44^+^ T cells, were more abundant in germ-free CD8.4 mouse than in germ-free CD8WT mouse (Fig. 2F-G, Fig. S2B). Collectively, these data implied that CD122^HI^ CD49d^-^ memory T cells represent the CD8^+^ VM T-cell population, which originates from naïve T cells with a relatively strong level of self-reactivity independently of foreign antigens.

### Virtual memory T cells use distinct TCR repertoire than naive T cells

Based on the previous data, generation of VM T cells can be understood as a fate decision checkpoint of individual naïve T cells (staying naïve vs. becoming VM), where the decision is based on the level of self-reactivity of the T cell’s TCR. This hypothesis predicts that naïve and VM T-cell compartments should contain different T-cell clones with distinct TCR repertoires. We analyzed the TCR repertoires by using a Vβ5 transgenic mouse with fixed TCRβ from the OT-I TCR [21]. The advantage of this mouse is that it generates a polyclonal population of T cells, but the variability between the clones is limited to TCRα chains. Moreover, pairing of TCRα and TCRβ upon repertoire analyses is not an issue in this model. Last, but not least, this mouse has a relatively high frequency of oligoclonal T cells that recognize the ovalbumin antigen [22].

Unimmunized Vβ5 mice have a frequency of memory CD8^+^ T cells around 10-15% which is comparable to wild type mice (Fig. 3A). When we analyzed the frequency of VM vs. naïve T cells among particular T-cell subsets defined by the expression of particular TCRVα segments, we observed that TCRVα3.2^+^ T cells are comparable to the overall population, TCRVα2^+^ T cells are slightly enriched for the VM T cells, and TCRVα8.3^+^ T cells have lower frequency of VM T cells than the overall population (Fig. 3A). These data suggested that naïve and VM T cells might have distinct TCR repertoires. However, the differences between the subsets were only minor, most likely because particular TCRVα subsets had still significant intrinsic diversity. To further reduce clonality in our groups, we gated on K^b^-OVA specific TCRVα3.2^+^, TCRVα2^+^, and TCRVα8.3^+^ T cells (Fig. 3B). Interestingly, around 50 % of OVA-specific TCRVα2^+^ clones exhibited VM phenotype, whereas vast majority of OVA-specific TCRVα8.3^+^ T cells were naïve and OVA-specific TCRVα3.2^+^ T cells had intermediate frequency of VM T cells in peripheral LN, mesenteric LN as well as in the spleen (Fig. 3B-C). Accordingly, the frequency of TCRVα2^+^ T cells is almost 10-fold higher in VM than in naïve OVA-specific T-cell population, whereas the frequency of TCRVα8.3^+^ T cells is slightly lower in VM than in naïve OVA-specific T cells (Fig. S3A). Similar differences between total and OVA-specific TCRVα3.2^+^, TCRVα2^+^, and TCRVα8.3^+^ CD8^+^ T cell subsets were observed in germ-free Vβ5 mice (Fig. 3D-E), confirming the VM identity of memory-phenotype T cells in the Vβ5 mice. In addition, OVA-specific TCRVα2^+^ had higher levels of CD5 than TCRVα8.3^+^ (Fig. S3B-C), suggesting that OVA-specific TCRVα2^+^ clones are on average more self-reactive than TCRVα8.3^+^ clones. This explains why more OVA-specific TCRVα2^+^ T cells than TCRVα8.3^+^ T cells acquire the VM phenotype in Vβ5 mice.

**Figure 3.**
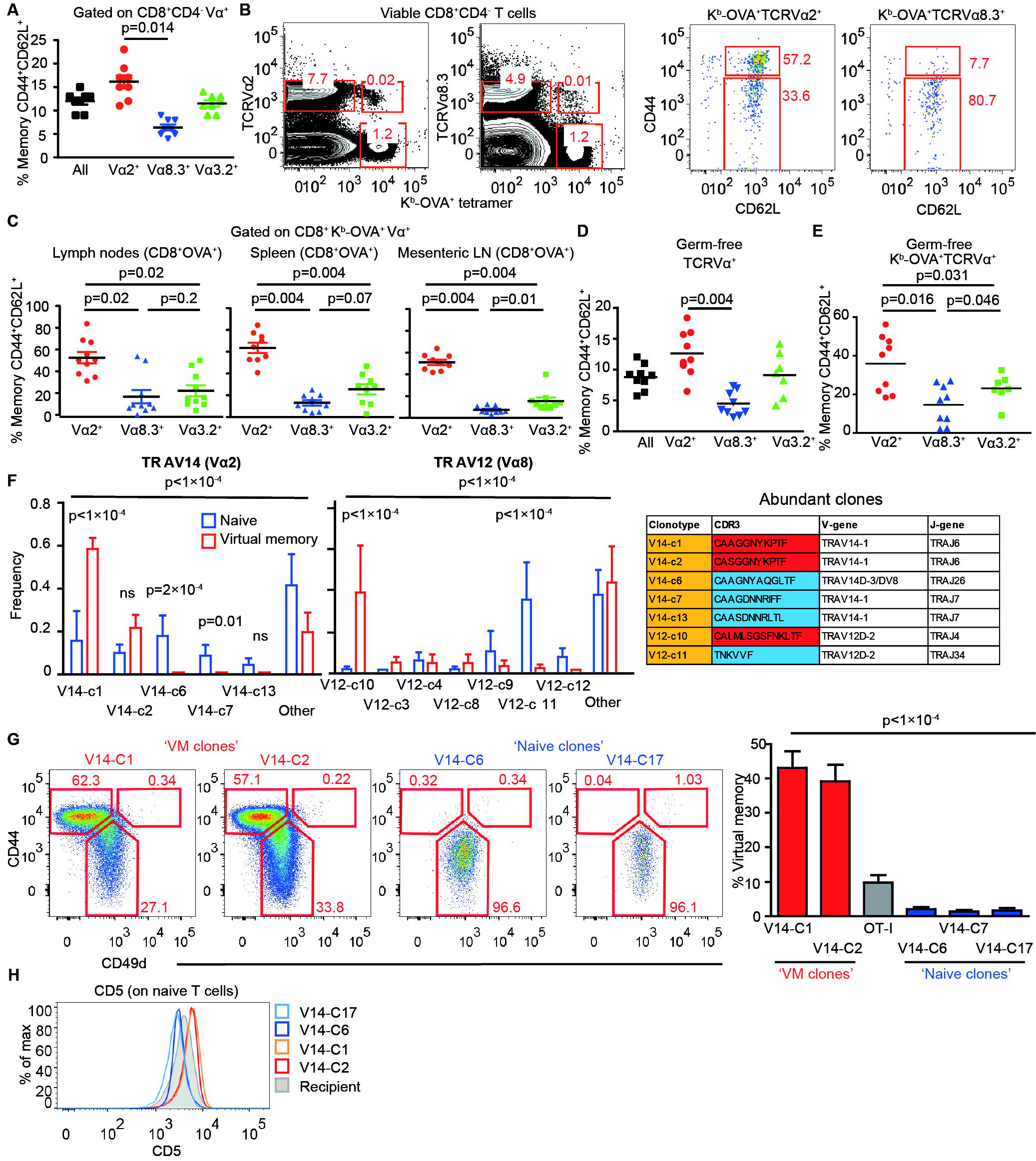
Virtual memory and naïve T cells use different TCR repertoires. (A) LN cells isolated from Vβ5 mice were stained for CD8, CD4, CD44, CD62L and Vα2 or Vα8.3 or Vα3.2. CD8^+^ T cells were gated as CD8^+^ CD4^-^ and then the percentage of CD44^+^ CD62L^+^ memory T cells among CD8^+^ Vα2^+^ or CD8^+^ Vα8.3^+^ or CD8^+^ Vα3.2^+^ T cells was determined by flow cytometry. Mean, n=8 mice from 8 independent experiments. (B-C) Cells isolated from peripheral LN (B-C), mesenteric LN (C) and the spleen (C) were stained as in (A) with the addition of OVA-tetramer. The OVA-reactive Vα-specific CD8^+^ T cells were gated and the percentage of CD44^+^ CD62L^+^ memory T cells was determined by flow cytometry. n=9-10 mice from. (D) The same experiment as in (A) was performed using germ-free Vβ5 mice. Mean, n=7-9 mice from 4 independent experiments. (E) The same experiment as in (B-C) was performed using a mixture of T cells isolated from LNs and the spleen from germ-free Vβ5 mice. Mean, n=7-9 from 2-3 independent experiments. (C-E) Statistical significance was determined by 2-tailed Wilcoxon signed-rank test. (F) RNA was isolated from memory (CD44^+^CD62L^+^) and (CD44^+^CD62L^+^) K^b^-OVA^+^ 4mer^+^ T cells sorted from LNs and the spleen of germ-free Vβ5 mice. TCRα encoding genes using either TRAV12 (corresponding to Vα8) or TRAV14 (corresponding to Vα2) were cloned and sequenced. 12-20 clones were sequenced in each group/experiment. Clonotypes identified in at least 2 experiments are shown. Mean frequency + SEM, n=4 independent experiments. Statistical significance was determined by Chi-square test (global test) and paired two-tailed T tests (individual clones). CDR3 sequences of clonotypes enriched in naïve or VM compartments are shown in the table. (G-H) Retroviral vectors encoding selected TCRα clones were transduced into immortalized Rag2^−/−^ Vβ5 bone marrow stem cells. These cells were transplanted into an irradiated Ly5.1 recipient. (G) At least 8 days after the transplantation, frequency of virtual memory T cells among LN donor T cells (CD45.2^+^ CD45.1^−^ GFP^+^) was analyzed. Mean+SEM; n=10-21 mice from 2-7 independent experiments. Statistical significance was tested using Kruskal-Wallis test. (H) CD5 levels on naïve monoclonal T cells was detected by flow cytometry. Representative mice out of 9-14 in total from 2-4 independent experiments. TM, true CM T cells.

Based on the analysis of particular TCRVα subsets, we hypothesized that naïve and VM T cells would contain different TCR clonotypes. We cloned and sequenced genes encoding for TCRα chains from OVA-reactive CD8^+^ VM and naïve T-cell subsets from germ-free Vβ5 mice using primers specific for TRAV14 (TCRVα2) and TRAV12 (TCRVα8) TCR genes (Table S1). The distribution of the clonotypes as well as TRAJ usage was significantly different between VM and naïve subsets (Fig. 3F, Fig. S3D). We observed essentially two types of clonotypes that were captured in more than 1 experiment (Fig. 3F). One type of clonotypes, called ‘VM clones’, was enriched among VM T cells and was also present in naïve T cells. The other type, ‘naïve clones’, was almost exclusively detected in naïve T cells. These data demonstrate that naïve and VM T-cell population contain different T-cell clones.

To directly investigate whether TCR is the main factor that determines whether a particular T cell has the potential to differentiate into VM T cells, we established a protocol to generate monoclonal T-cell populations using transduction of a particular TCRα-encoding gene in a retrogenic vector into immortalized hematopoietic stem cells [23]. We transduced immortalized Vβ5 Rag2^−/−^ hematopoietic stem cells with expression vectors encoding for 2 VM TCRα clones (V14-C1 and V14-C2), 3 naïve TCRα clones (V14-6, V14-7, and V14-17), or OT-I TCRα as a control (Fig. S3E). At least 8 weeks after the transplantation of the progenitors into a Ly5.1 recipient, we analyzed the cell fate of the donor monoclonal T-cell populations. T cells expressing VM TCR clones formed a significant VM T-cell population, whereas T cells expressing naïve TCR clones formed a homogenous naïve population (Fig. 3G, Fig. S3F). These data demonstrate that the virtual memory T cells are formed only from certain T-cell clones and that the TCR sequence determines whether a T cell differentiates into a VM T cell or stays naïve.

In a next step, we addressed whether ‘VM clones’ are more self-reactive than ‘naïve clones’. We compared levels of CD5, a commonly used reporter for self-reactivity, on naive (CD44^’^) populations of retrogenic monoclonal T cells. ‘VM clones’ expressed significantly higher CD5 levels than ‘naïve clones’, indicating that ‘VM clones’ are indeed T-cells with a relatively high level of self-reactivity (Fig. 3H, Fig. S3G). Interestingly, retrogenic OT-I T cells represented an intermediate VM population, which corresponds to their level of self-reactivity (Fig. 1H, Fig. 3G, Fig. S3F-G). Moreover, the relative size of retrogenic OT-I VM population well corresponded to the frequency of VM T cells in conventional transgenic OT-I TCR mice (Fig. 1E, H), indicating that the protocol for generation of retrogenic monoclonal T cells itself does not have a strong effect on VM formation. Overall, these results document that only relatively highly self-reactive clones form VM T cells.

### Virtual memory T cells represent an intermediate stage between naïve and memory T cells

The relationship of the differentiation programs of naïve, VM, and true CM T cells is unclear. So far, CD49d and, to a lesser extent, CD122 were the only known markers discriminating between true CM memory and VM T cells [2, 4, 5, 9]. To compare their gene expression profiles, we performed deep RNA sequencing of the transcripts from sorted CD8^+^ naïve (CD44^−^ CD62L^+^) and VM (CD44^+^ CD62L^+^ CD49d^−^) T cells isolated from germ-free animals and from true antigen-specific CM T cells (K^b^-OVA+ CD44^+^ CD62L^+^), generated by infecting Vβ5 mice with Lm expressing OVA (Lm-OVA). The data showed that naïve, VM, a true CM T cells represent three distinct T cell populations (Fig. S4A-B). We identified genes differently expressed in VM T cells in comparison to naïve or true CM T cells (Tables S2-5). Based on previously published data [24–26], we established lists of memory signature and naïve signature genes. As expected, memory signature genes were enriched in true CM T cells and naïve signature genes were enriched in naïve T cells (Fig. 4A, S4C). Interestingly, VM T cells exhibited an intermediate gene expression profile (Fig. 4A, S5C). Pairwise rotation gene set tests revealed the hierarchy in the enrichment for memory signature genes and for naïve signature genes as true CM > VM > naïve, and naïve > VM > true CM, respectively (Fig. 4B). VM T cells also showed an intermediate expression of cytokine and chemokine encoding genes (Fig. 4C, Fig S4D). The transcription of genes encoding for cytokine and chemokine receptors in VM T cells seemed to be also somewhere half-way between naïve and true CM T cells (Fig. S4E-F).

**Figure 4.**
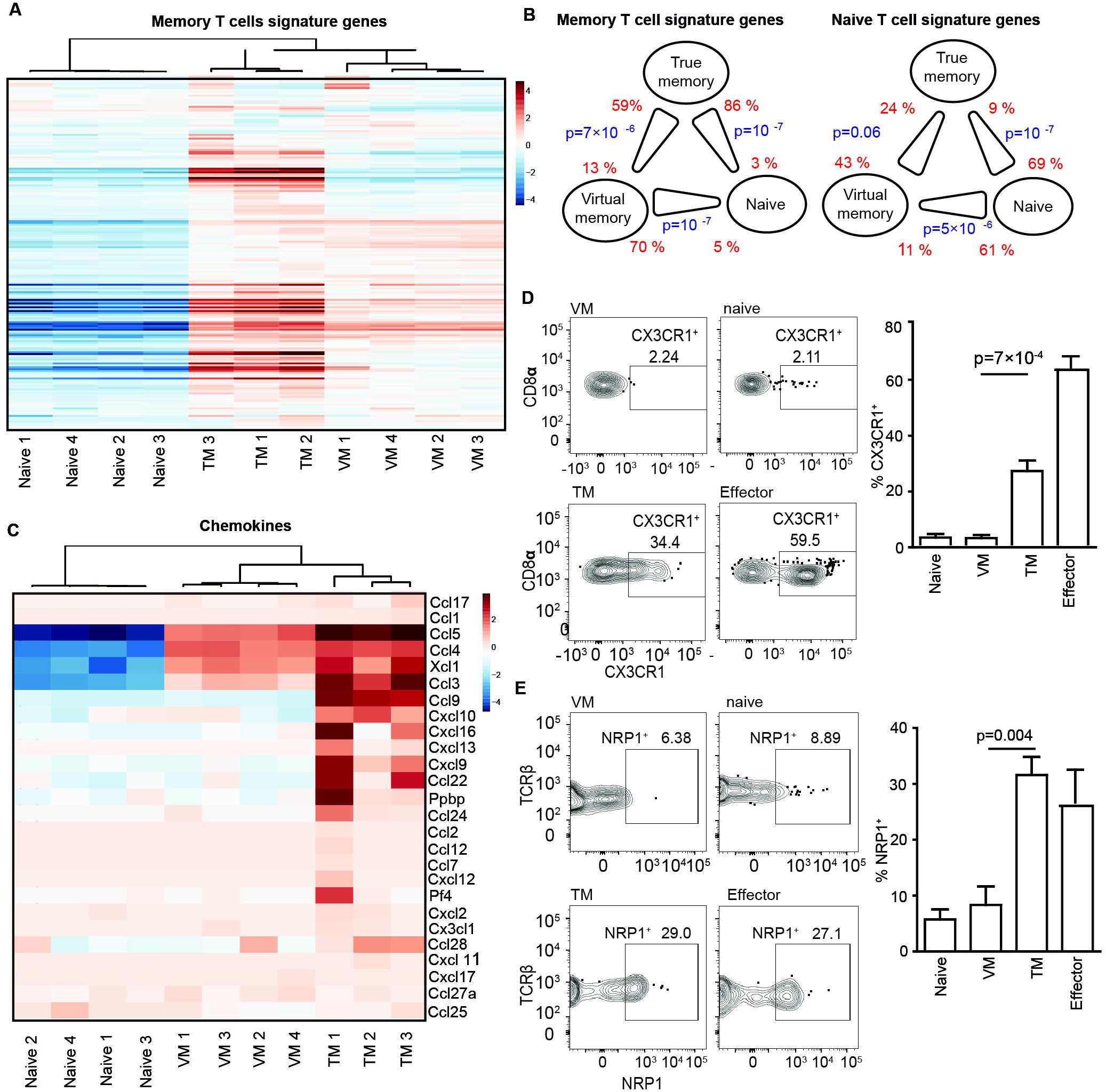
Virtual memory T cells represent an intermediate stage between naïve and true memory T cells. (A-C) Transcriptomes of naive (n=4), VM (n=4), and true CM (n=3) CD8^+^ T cells were analyzed by deep RNA sequencing. (A) Enrichment of CD8^+^ memory signature genes (as revealed by previous studies) in naïve, virtual memory, and true memory T cells. (B) Pairwise comparisons between naïve, VM, and true CM CD8^+^ cells for the overall enrichment of the memory signature and naïve signature gene sets by a method ROAST. The thick end of the connecting line between the populations indicates the population with the overall relative enrichment of the gene set, the percentage of the genes from the gene set that are more expressed in the indicated population (z-score > sqrt(2)) over the opposite population is indicated. (C) The relative enrichment of chemokine encoding transcripts in the samples is shown. (D-E) Surface staining for CX3CR1 (D) and NRP1 (E) was performed on naïve (gated as CD62L^+^CD44^−^CD49d^low^) and VM (CD62L^+^CD44^+^CD49d^low^) K^b^-OVA-4mer^+^ CD8^+^ T cells isolated from unprimed Vβ5 mouse and on true CM memory (gated as CD62L^+^CD44^+^CD49d^high^) and effector/effector memory (CD62L^−^CD44^+^CD49d^high^) K^b^-OVA-4mer^+^ CD8^+^ T cells isolated from Vβ5 mouse 30-45 days after Lm-OVA infection. A representative experiment out of 5 (D) or 4 (E) in total is shown. Mean percentage + SEM of CX3CR1^+^ and NRP1^+^ cells within the particular population is shown. (D) n=10 immunized mice and 5 unprimed mice from 5 independent experiments. (E) n=8 immunized mice and 4 unprimed mice from 4 independent experiments. Statistical analysis was performed by 2-tailed Mann-Whitney test.

We further investigated selected differentially expressed genes between VM and true CM cells on a protein level. RNA encoding for CX3CR1 and NRP1 showed enrichment in true CM vs. VM T cells. For this reason, we compared surface levels of CX3CR1 and NRP1 on naïve and VM T cells from unprimed mice and on true CM and effector/effector memory T cells from LM-OVA infected mouse during the memory phase (Fig. 4D-E). In contrast to VM T cells, a significant percentage of effector and true CM T cells expressed CX3CR1 and NRP1, confirming the transcriptomic data and suggesting that VM T cells can be characterized as CX3CR1 and NRP1 negative.

### Differentiation into virtual memory T cells does not break self-tolerance

VM T cells provide better protection against Lm than naïve T cells and respond to proinflammatory cytokines IL-12 and IL-18 by producing IFNγ [2, 5, 10]. Moreover, VM T cells express higher levels of several killer lectin-like receptors than naïve T cells (Table S4 and [5]). The combination of a hyperresponsive differentiation state with the expression of highly self-reactive TCRs suggests that VM CD8^+^ T cells could be less self-tolerant than naïve T cells and might represent a risk for inducing autoimmunity. We used CD8WT OT-I T cells (mostly naïve) and CD8.4 OT-I T cells (mostly VM) for a functional comparison of naïve and VM T cells with the same TCR specificity. Because CD8.4 T cells have stronger reactivity to antigens than CD8WT T cells, this monoclonal T-cell model corresponds to the physiological situation where TCRs of VM T cells are intrinsically more reactive to self-antigens than TCRs of naïve T cells. VM T cells, but not naïve OT-I T cells, rapidly produced IFNγ after the stimulation by PMA/ionomycine or cognate antigen (Fig. 5A), supporting the idea of an autoimmune potential of VM T cells. We directly tested this hypothesis using an experimental model of type I diabetes [27]. We transferred CD8WT OT-I or CD8.4 OT-I T cells into RIP.OVA mice expressing OVA under the control of rat insulin promoter [28] and primed them with Lm-OVA or Lm-Q4H7 [27]. Q4H7 is an antigen that binds to the OT-I TCR with a low affinity and does not negatively select OT-I T cells in the thymus [14, 29]. Thus, Q4H7 resembles a self-antigen that
positively selected peripheral T cells might encounter at the periphery. Surprisingly, CD8.4 OT-I T cells were not more efficient in inducing the autoimmune diabetes than CD8WT OT-I T cells in any tested condition (Fig. 5B-C). Accordingly, CD8WT and CD8.4 OT-I T cells show comparable expansion when primed by Lm-OVA or Lm-Q4H7 in vivo (Fig. S5A-B).

**Figure 5.**
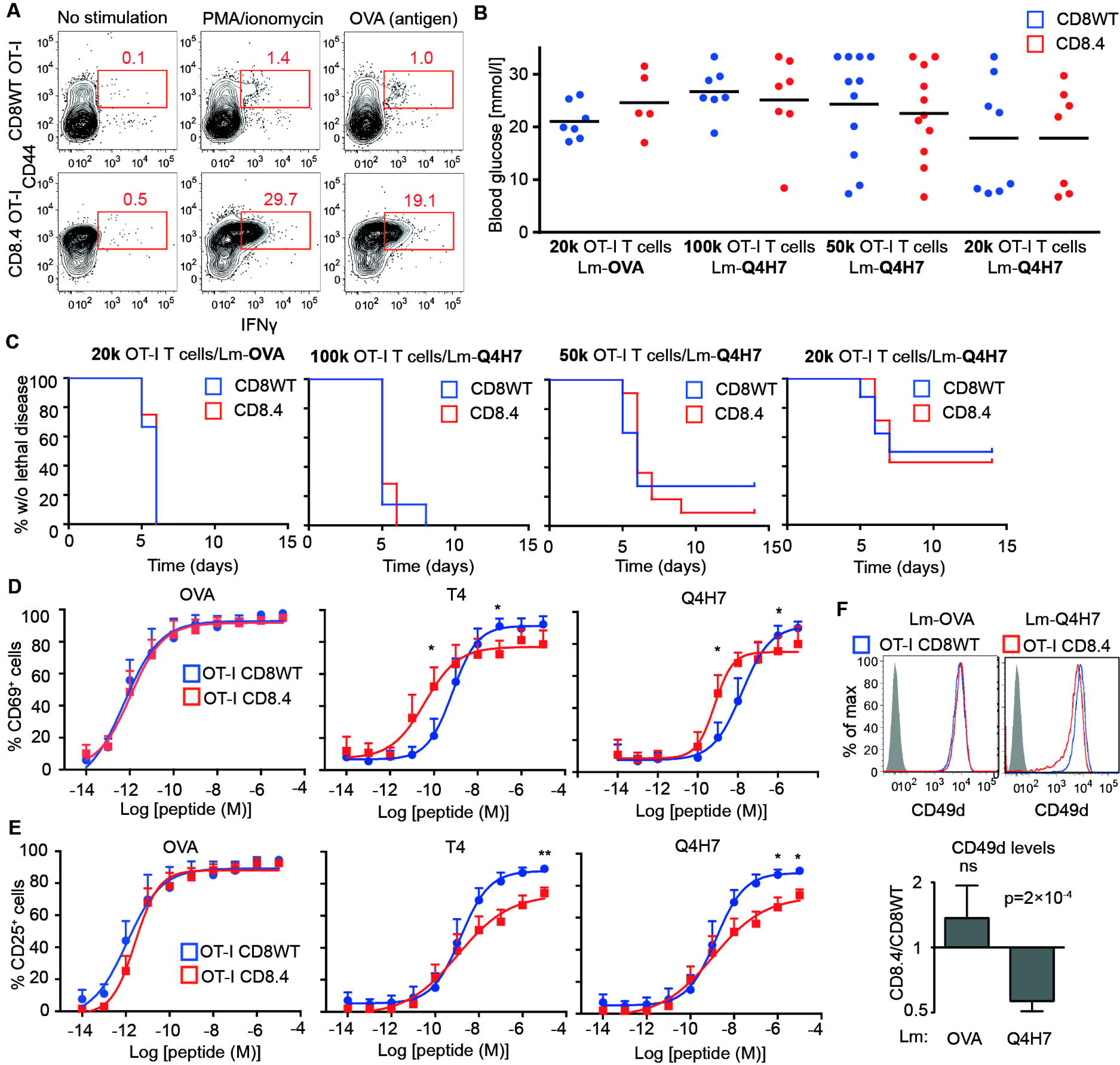
Virtual memory T cells are as tolerant to low affinity antigens as naïve T cells. (A) LN cells isolated from CD8WT OT-I and CD8.4 OT-I mice were stimulated with PMA and ionomycine or OVA peptide in the presence of BD Golgi Stop for 5 hours and the production of IFNγ was analyzed by flow cytometry (gated as CD8^+^). A representative experiment out of 4 in total. (B-C) Indicated number of CD8WT OT-I or CD8.4 OT-I T cells were adoptively transferred into RIP-OVA hosts, which were infected with Lm-OVA or-Q4H7 one day later. (B) Blood glucose was measured 7 days after the infection. (C) The glucose in the urine was monitored for 14 days. The percentage of non-diabetic mice in time is shown. Differences between OTI-I and CD8.4 were not significant by Log-Rank test. n=5-11 mice in 3-6 independent experiments. (D-E) CD8WT OT-I or CD8.4 OT-I LN T cells were stimulated ex vivo with dendritic cells loaded with varying concentrations of OVA, Q4R7, Q4H7 peptides overnight and the expression of CD69 (D) and CD25 (E) on CD8^+^ T cells was analyzed. Mean + SEM. n=3-4 independent experiments. Statistical significance was determined paired 2-tailed Student’s T test (C, D), and one value T test (E). * p < 0.05, **<0.01 (F) A CD8WT OT-I or CD8.4 OT-I LN T cells were adoptively transferred to polyclonal Ly5.1 host mice, which were infected one day later with transgenic Lm-OVA or-Q4H7. 6 days after the infection, splenocytes from the hosts were isolated and analyzed for CD49d expression. Statistical significance was determined using a one-value two-tailed T test (for the ratio of CD49d MFI between the subsets).

To further analyze the functional responses of VM T cells, we stimulated CD8.4 OT-I and C8WT OT-I T cells with dendritic cells loaded with OVA or suboptimal cognate antigens T4 or Q4H7 ex vivo. No significant difference in the upregulation of CD69 or CD25 between naïve and VM cells was observed in the case of high-affinity OVA stimulation. However, the responses to antigens with suboptimal affinity to the TCR differed between these two cell types. Interestingly, although, CD8.4 OT-I T cells showed stronger CD69 upregulation than naïve T cells when stimulated with low antigen dose, when the antigen dose was high, the response of VM T cells was lower than that of naïve T cells (Fig. 5D). Upregulation of CD25 was lower in CD8.4 OT-I T cells than in naïve T cells, when activated with the low-affinity antigens (Fig. 5D-E). Moreover, CD8.4 OT-I T cells did not upregulate CD49d (a subunit of VLA-4) to the same extent as CD8WT OT-I upon immunization with Lm-Q4H7 (Fig. 5F). Overall, although we and others showed that VM T cells elicit stronger responses than naïve T cells in some assays (IFNγ production, protection against Lm), they do not show a higher autoimmune potential than naïve T cells, most likely because they possess additional mechanisms to suppress their effector functions. One such mechanism, contributing to the self-tolerance of VM T cells, can be lower expression of CD49d and CD25 upon activation with suboptimal antigens. Altogether, these data establish that relatively strongly self-reactive T-cell clones differentiate to VM T cells and trigger a specific developmental program that enables them to efficiently response to infection, but does not increase their autoimmune potential (Fig. 6).

**Figure 6.**
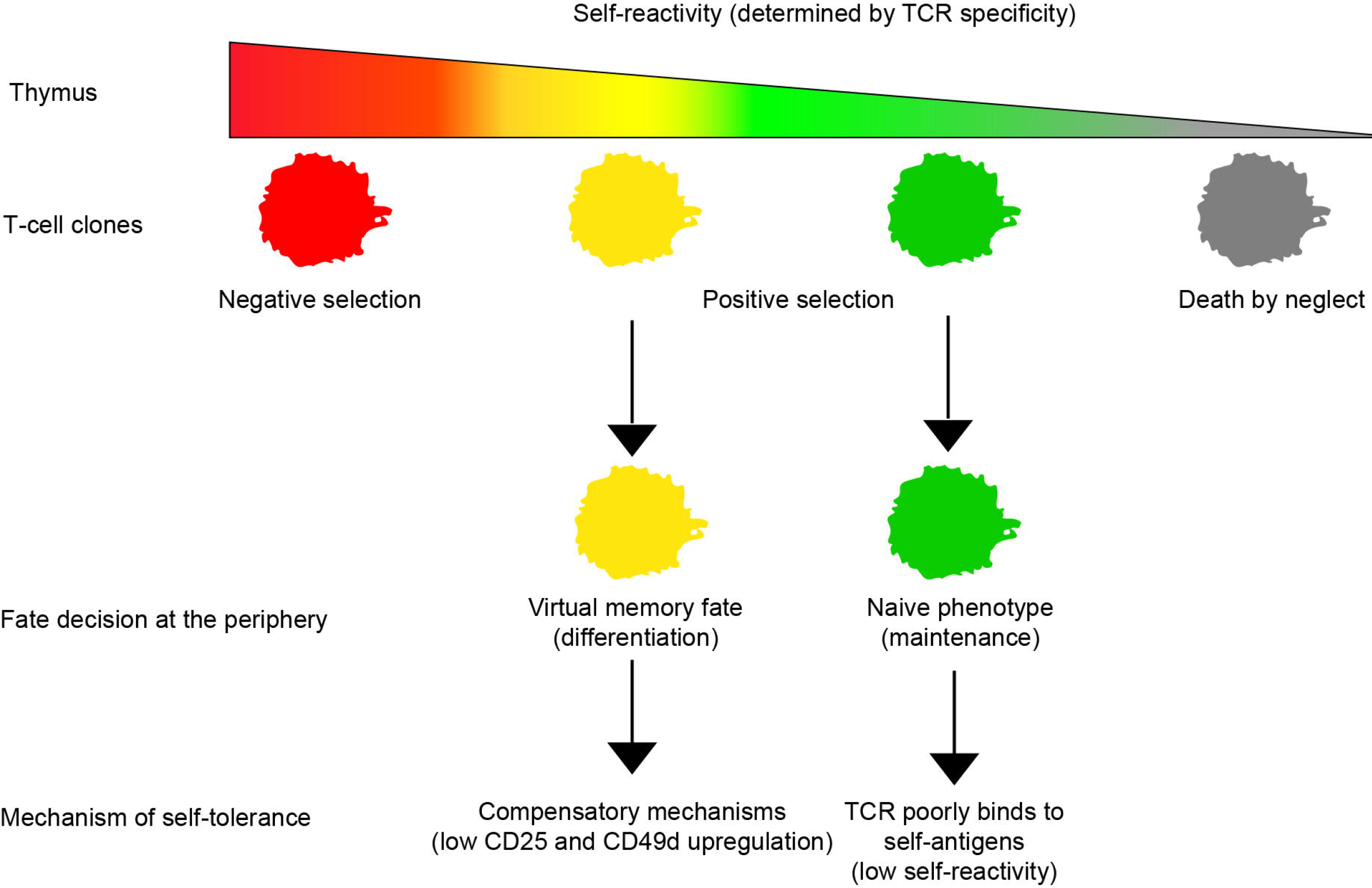
Schematic representation of the role of self-reactivity in major cell fate decisions of conventional CD8^+^ T cells. Our results establish a novel T-cell fate decision checkpoint, differentiation of positively selected T-cell clone with a relatively high level of self-reactivity into virtual memory T cells.

## Discussion

We observed that CD8-Lck coupling frequency regulates intensity of TCR homeostatic signals. For the first time, we showed that the intrinsic sensitivity of the TCR signaling machinery sets the frequency of VM CD8^+^ T cells. We also showed that only relatively strongly self-reactive T-cell clones have the potential to form VM T cells. We identified the gene expression profile of VM T cells and showed that they represent an intermediate stage between naïve and true CM T cells. Although the combination of relatively strong self-reactivity and acquisition of the partial memory program could represent a potential risk for autoimmunity, we observed that VM T cells are not more potent than naïve T cells in a model of experimental type I diabetes.

It is well established that some memory-phenotype T cells respond to an antigen, they have not been previously exposed to [2–4, 10, 30]. Some researchers call these cells VM T cells and propose that they were generated in the absence of a foreign antigenic stimulation [2, 30]. The main argument supporting this hypothesis is that germ-free mice, with low levels of foreign antigenic exposure, contain comparable levels of CM-phenotype T cells as control mice [2]. However, in mice with normal microbiota, which were used for the subsequent characterization of VM T cells, it is difficult to exclude the existence of cross-reactive memory T cells that were previously exposed to another foreign antigen [3]. Importantly, we show that CD8.4 knock-in mice with enhanced homeostatic TCR signaling exhibit larger VM compartment than CD8WT T cells in SPF and germ-free conditions. We confirmed that VM T cells have lower expression of CD49d than antigen-experienced cells using germ-free mice and confirmed that antigen-inexperienced VM T cells can be defined as CD49^−^ CD122^HI^ T cells. Altogether these data established that the strength of homeostatic signals provided to T cells is a major factor leading to formation of VM compartment independently of stimulation with cognate foreign antigens.

We demonstrated that, in contrast to VM T cells, the population of putative ‘antigen-experienced’ CD8^+^ CD44^+^ CD49d^+^ T cells is not regulated by the magnitude of the homeostatic TCR signaling. Because the size of CD122^LOW^ CD49d^+^ CD8^+^ T-cell subset is comparable between germ-free and SPF mice, the identity and origin of these cells in unimmunized mice is enigmatic. It is not clear whether this T-cell subset is induced by residual environmental antigens in the germ-free mice or contains a homeostatic memory-phenotype cell population unrelated to VM T cells.

It has been observed that IL-15 availability is a limiting factor regulating the size of the VM subset [4, 5]. In this study, we showed that the intrinsic sensitivity of the TCR signaling machinery (specifically the CD8-Lck coupling frequency) is another major factor that sets the frequency of VM CD8^+^ T cells in the secondary lymphoid organs. It has been suggested that the level of CD5, a marker of self-reactivity, is linked with the T-cell ability to form VM T cells [5]. We investigated individual T-cell clones using transgenic cells with normal and hypersensitive TCR signaling machinery, comparing TCR repertoires of naïve and VM T cells, and analyzing retrogenic monoclonal T-cell populations. These complementary approaches revealed that VM T-cell formation absolutely depends on the level of self-reactivity of a particular T cell and exhibits a threshold behavior. Relatively highly self-reactive T cell clones frequently differentiate into VM T cells (~40-50%), whereas weakly self-reactive T cells completely lack this property. This finding characterizes the formation of VM T cells as a previously unappreciated T-cell fate decision check-point, where the intensity of homeostatic TCR signals is the critical decisive factor. Our data also explains a previous observation that VM T cells are formed exclusively from T cells expressing endogenous recombined TCR chains in OT-I Rag^+^ mice during aging [13]. Some of the T-cell clones that replaced the OT-I TCR with a variable endogenous one are probably more self-reactive than OT-I T cells, which drives their differentiation in VM T cells.

The functionality of a T cell subset is determined by its gene expression profile. Whereas it is clear that VM T cells substantially differ from naïve T cells [2, 4, 5, 10], the CD49d and CD122 were the only markers that can distinguish VM T cells from true CM T cells. In this study, we characterized gene expression of VM T cells and compared it to naïve T cells from germ-free mice and foreign antigen-specific true CM T cells. Analysis focusing on previously established naïve and memory T cell signature genes revealed that VM T cells have an intermediate gene expression profile between naïve and true CM T cells. Accordingly, expression of chemokines and cytokines was generally lower in VM T cells than in true CM T cells. These data suggest that VM T cells trigger a partial memory program. Alternatively, true CM CD8^+^ T cells might represent a heterogeneous population of two or more subsets with different degrees of similarity to VM T cells, as suggested by heterogeneous expression of CX3CR1 and NRP1, two genes that showed a large difference between true CM and VM T cells. Indeed, CX3CR1 has been proposed as a marker that discriminates different subsets of true memory T cells [31, 32]. Single cell gene expression profiling would show whether immune responses to foreign antigens generate any true memory T cells identical to VM T cells.

Based on pilot studies in the field [33–35], it was generally accepted that one feature of immunological memory is that a memory T cell elicits a faster and stronger response to cognate antigens than a naïve T cell. However, recent evidence showed that, at least under certain conditions, the response of naïve T cells to an antigen is stronger than the response of memory T cells [36–38]. VM T cells were previously shown to surpass naïve T cells in their response to inflammatory cytokines IL-12 and IL-18 [2], in the rapid generation of short-lived effectors [10], and in the protection against Lm in both antigen-specific [10] and by-stander manners [39]. These findings, showing potent effector potential of VM T cells, together with our data, demonstrating that VM T cells develop from relatively strongly self-reactive T-cell clones, indicate that VM T cells might represent a high risk for breaking self-tolerance and inducing autoimmunity. We addressed this issue by using CD8WT OT-I Rag2^−/−^ and CD8.4 OT-I Rag2^−/−^ T cells as monoclonal models for naïve and VM T cells, respectively, taking advantage of the fact that these T cells express the same TCR and have the same antigenic specificity. Moreover, this setup mimics the enhanced response of VM T cells to self-antigens as well as the differences based on the naïve or VM differentiation stages. We observed that VM T cells, but not naïve T cells, rapidly secrete IFNγ after antigenic stimulation. However, VM T cells were not more efficient than naïve T cells in inducing the experimental autoimmune diabetes. This can be at least partially explained by the fact that VM T cells show lower upregulation of CD25 and VLA-4 than naïve T cells when activated with a suboptimal antigen. VLA-4 has been previously shown to be essential for the induction of the tissue pathology in the mouse experimental model of type I diabetes [27]. We propose that VM T cells surpass naïve T cells in some kind of responses to promote rapid immunity to pathogens [5, 10], but they also develop compensatory mechanisms that make these cells self-tolerant to an extent comparable to naïve T cells. We used an experimental model of type I diabetes, which is a prototypic autoimmunity that involves self-reactive CD8^+^ T cells. We cannot exclude the possibility that VM T cells represent a major risk in other types of autoimmune diseases/conditions. However, our data from monoclonal models for VM and naïve CD8^+^ T cells have general implications that comparison of particular T-cell subsets highly depends on the immunological context (e.g., protective immunity vs. autoimmunity). Given the controversy concerning the reactivity of naïve and true CM T cells, it is possible that similar ambiguity (i.e., hyperresponsiveness in some aspects, while hyporesponsiveness in others) applies to true memory T cells as well.

Recently, it has been proposed that human innate Eomes^+^ KIR/NKG2A^+^ CD8^+^ T cells [40] represent human counterparts of murine VM T cells, although expression of some markers including CD27 and CD5 substantially differed between these two subsets [5]. It would be interesting to elucidate whether the human Eomes^+^ KIR/NKG2A^+^ CD8^+^ subset shows similar gene expression pattern as mouse VM T cells, whether they originate from relatively highly self-reactive clones, and whether these cells acquire tolerance mechanisms as murine VM CD8^+^ T cells do.

## Author Contribution

AD, AM, DM, MH, and OS planned, performed, and analyzed experiments. VH and MP performed and analyzed experiments. PD analyzed experiments. RI did the bioinformatic analysis of the RNA sequencing data. SO and DZ prepared transgenic Lm strains and provided Vβ5 mice. KDM established and managed the germ-free strains. OS conceived the study. OS, PD, and AM wrote the manuscript. All authors commented on the manuscript.

## Acknowledgement

We thank Prof. Ed Palmer for his multisided support to the project. We thank Prof. Alfred Signer for providing us with CD8.4 mice. We thank Ladislav Cupak, Barbara Hausmann, and Rosmarie Lang for technical assistance and genotyping of mice. This project was supported by the Swiss National Science Foundation (Promys, IZ11Z0_166538), the Czech Science Foundation (GJ16-09208Y) to OS and ERC Starting Grant (ProtecTC) to DZ. The Group of Adaptive Immunity at the Institute of Molecular Genetics in Prague is supported by an EMBO Installation grant. AM, VH, and MP are students of the Faculty of Science, Charles University, Prague.

## Material and Methods

### Antibodies and reagents

Antibodies to following antigens were used for flow cytometry: CD69 (clone H1.2F3), CD11a (LFA-1) (clone M17/4), CD25 (PC61), CD3 (145-2C11), IFNγ (XMG1.2), TCRβ (H57-597), TCR Vα2 (B20.1), TCR Vα8.3 (B21.14), TCR Vα3.2 (R3-16), CD49d (R1-2), CD5 (53-7.3) (all BD Biosciences), Tbet (4B10), Eomes (Dan11mag), CD8α (53-6.7), CD8β (H35-17.2), CD127 (A7R34) (all eBioscience) CD44 (IM7), CD4 (RM-45), CD62L (MEL-14), CD122 (TM-beta1), KLRG1(2F1), PD-1 (RMP1-30), CD19 (6D5) (all Biolegend), pErk1/2 (D13.14.4E, Cell Signaling). The antibodies were conjugated with various fluorescent dyes or with biotin by manufacturers. K^b^-OVA PE tetramer was prepared as described previously [14]. Peptides OVA (SIINFEKL), Q4R7 (SIIRFERL), Q4H7 (SIIRFEHL), NP68 (ASNENMDAM), and NP372E (ASNENMEAM) were purchased from Eurogentec or Peptides&Elephants.

### Flow cytometry and cell counting

For the surface staining, cells were incubated with diluted antibodies in PBS/0.5% gelatin or PBS/2% goat serum on ice. LIVE/DEAD near-IR dye (Life Technologies) or Hoechst 33258 (Life Technologies) were used for discrimination of live and dead cells. For the intracellular staining, cells were fixed and permeabilized using Foxp3/Transcription Factor Staining Buffer Set (eBioscience, 00-5523-00). For some experiments, enrichment of CD8^+^ T cells was performed using magnetic bead separation kits EasySep (STEMCELL Technologies) or Dynabeads (Thermo Fisher Scientific) according to manufacuter’s instructions prior to the analysis or sorting by flow cytometry. Cells were counted using Z2 Coulter Counter (Beckman) or using AccuCheck counting beads (Thermo Fisher Scientific) and a flow cytometer. Flow cytometry was carried out with a FACSCantoII, LSRII or a LSRFortessa (BD Bioscience). Cell sorting was performed using a FACSAria III or Influx (BD Bioscience). Data were analyzed using FlowJo software (TreeStar).

### Experimental animals

All mice were 5-12 weeks old and had C57Bl/6j background. RIP.OVA [28], OT-I Rag2^−/−^ [41], CD8.4, F5 Rag1^-/-^ [15], and Vβ5 [21] strains were described previously. Mice were bred in our facilities (SPF mice: University Hospital Basel and Institute of Molecular Genetics; germ-free mice: University of Bern, Switzerland) in accordance with Cantonal and Federal laws of Switzerland and the Czech Republic. Animal protocols were approved by the Cantonal Veterinary Office of Baselstadt, Switzerland, and Czech Academy of Sciences, Czech Republic. Transfer into germ-free conditions was performed using time-mating followed by transferring 2-cell embryos into germ-free foster mothers.

Males and females were used for the experiments. If possible, age-and sex-matched pairs of animals were used in the experimental groups. If possible, littermates were equally divided into the experimental groups. No randomization was performed. The experiments were not blinded since no subjective scoring method was used.

### Ex vivo activation assay

For the analysis of IFNγ production, T cells (1×10^6^/ml in RPMI/10% FCS) were stimulated with 10 ng/ml PMA and 1.5 μM ionomycine or 1 μM OVA peptide in the presence of BD Golgi Stop for 5 hours. For the CD69 and CD25 upregulation assay, dendritic cells differentiated from fresh or immortalized bone marrow stem cells were pulsed with indicated concentration of indicated peptides and mixed with T cells isolated from LNs in a 1:2 ratio as described previously [41].

### RNA sequencing

RNA was isolated using Trizol (Thermo Fisher Scientific) followed by in-column DNAse treatment using RNA clean&concentrator kit (Zymo Research). The library preparation and RNA sequencing by HiSeq2500 (HiSeq SBS Kit v4, Illumina) was performed by the Genomic Facility of D-BSSE ETH Zurich in Basel. Obtained single-end RNA-seq reads were mapped to the mouse genome assembly, version mm9, with RNA-STAR [42], with default parameters except for allowing only unique hits to genome (outFilterMultimapNmax=1) and filtering reads without evidence in spliced junction table (outFilterType="BySJout"). All subsequent gene expression data analysis was done within the R software (R Foundation for Statistical Computing, Vienna, Austria). Raw reads and mapping quality was assessed by the qQCreport function from the R/Bioconductor software package QuasR (version 1.12.0, [43]). Using RefSeq mRNA coordinates from UCSC (genome.ucsc.edu, downloaded in July 2013) and the qCount function from QuasR package, we quantified gene expression as the number of reads that started within any annotated exon of a gene. The data are deposited in the GEO database (GSE90522). The differentially expressed genes were identified using the edgeR package (version 3.14.0, [44]). We generated lists of naïve and memory signature genes based on previously published studies (gene sets M3022, M5832, M3039 for memory, and M3020, M5831, M3038 for naïve T cells in the Molecular Signature Databases [45]) [24–26]. In our memory and naïve signature gene lists, we included only genes that were listed at least in 2 out of the above-mentioned 3 respective gene sets. For the global comparison of the expression of the signature genes in naïve, VM, and true CM T cells, we used self-contained gene set enrichment test called Roast, which is available in edgeR package [39].

### Bone marrow chimeras

Bone marrow cells were isolated from long bones of indicated mouse strains and lymphocytes were depleted using biotinylated antibodies to CD3 and CD19 and Dynabeads biotin binder kit (Thermo Fisher Scientific). In total 6×10^6^ cells (always a 1:1 mixture from 2 different donor strains) in 200 μL of PBS, were injected into irradiated (300 cGy) Rag2^−/−^ host mice i.v. The mice were analyzed 8 weeks after the transfer.

### DNA cloning and production and viruses

RNA was isolated using Trizol reagent (Thermo Fisher Scientific) and RNA clean&concentrator kit (Zymoresearch, R1013). Reverse transcription was performed using RevertAid (Thermo Fisher Scientific) according to the manufacturer’s instructions. TCR sequences were amplified using cDNA from sorted T cells by PCR using Phusion polymerase (New England Biolabs) and following primers: TRACrev (EcoRI) 5’-TCAGACgaattcTCAACTGGACCACAGCCTCA, TRAV14for (XhoI) 5’ GTAGCTctcgagATGGACAAGATCCTGACAGCA, TRAV12for (XhoI) 5’ GTAGCTctcgagATGCGTCCTGDCACCTGCTC and ligated into pBlueScript vector using T4 ligase (New England Biolabs) and sequenced by Sanger sequencing using T7 primer 5’ TAATACGACTCACTATAGGG. Selected clones were cloned into MSCV-GFP vector via EcoRI and XhoI (New England Biolabs).

Coding sequence of SCF, IL-3 and IL-6 was obtained from bone marrow cDNA with Phusion polymerase using these primers: SCFfor 5’-TTGGATCCGCCACCATGAAGAAGACACAAACTTGGATTATC, SCFrev 5’-AACTCGAGTTACACCTCTTGAAATTCTCTCTCTTTC, IL-3for 5’-TTGAATTCGCCACCATGGTTCTTGCCAGCTCTACCACCAG, IL3rev 5’-AACTCGAGTTAACATTCCACGGTTCCACGGTTAGG, IL-6for 5’-TTGAATTCGCCACCATGAAGTTCCTCTCTGCAAGAGACTT, IL6rev 5’ AACTCGAGCTAGGTTTGCCGAGTAGATCTCAAAGTG. cDNA was cloned into pXJ41 expression vector using BamHI or EcoRI and XhoI restriction sites and sequenced. Cytokines were produced in HEK293 cells transfected with pXJ41 using polyethylenimine (PEI) transfection in ratio 2.5 μl PEI to 1 μg DNA). Supernatant was harvested 3 days after transfection. Titration against commercial recombinant cytokines of known concentration and their effect on BM cells proliferation in vitro was used to determine biological activity of cytokines in supernatant. Dilution of supernatant corresponding to concentration of 100 ng/ml SCF, 20 ng/ml IL-3 and 10 ng/ml IL-6 was used for cultivation of immortalized bone marrow cells.

Retroviral MSCV and pMYs particles were generated by transfection of the vectors into Platinum-E cells (Cell Biolabs) by PEI as described above.

### Monoclonal retrogenic T cells

Generation of immortalized bone marrow was described previously [23]. Briefly, Vβ5 Rag2^−/−^ mice were treated with 100mg/kg 5-fluorouracil and bone marrow cells were harvested 5 day later. Cells were cultivated in complete IMDM supplemented with SCF, IL-3, and IL-6. After 2 days, proliferating cells were virally transduced with a fusion construct NUP98-HOXB4 in a retroviral vector pMYs. Viral infections were performed in the presence of 10 μg/ml polybrene by centrifugation (90 min, 1250g, 30 °C). The transduced cells were selected for 2 days in puromycine (1 μg/ml). Selected immortalized cells were subsequently virally transduced with MSCV vector containing s TCRα-encoding gene and GFP as a selection marker. Two days after the transduction, GFP^+^ were FACS sorted and transplanted into irradiated (7 Gy) congenic Ly5.1 recipients. At least 8 weeks after the transplantation, the recipient mice were sacrificed and donor LN T cells were used for cell fate analysis by flow cytometry.

### In vivo activation

Indicated numbers of T cells were adoptively transferred into a host mouse i.v. On a following day, the host mice were immunized with indicated peptide (50 μg) and LPS (25 μg) in 200 μL of PBS i.p. or with 5000 CFU of Lm. Lm strains expressing OVA, Q4R7, and Q4H7 have been described previously [27]. Lm expressing NP68 was produced by adding the ASNENMDAM epitope to ovalbumin encoding gene and introducted to Lm as previously described [46]. True CM T cells were generated by infecting Vβ5 mice with Lm-OVA. After 60-90 days (for RNAseq) or 30-50 days (for FACS staining), CD8^+^ Kb-OVA tetramer^+^ CD49d^+^ CD44^+^ CD62L^+^ from LNs and the spleen were sorted (or gated). In the experimental model of autoimmunity, we monitored glucose in the urine on a daily basis using test strips (Diapure-Test 5000, Roche). If not indicated, the animal was considered to suffer from lethal autoimmunity when the concentration of glucose in the urine reached ≥ 5000 mg/dL. We also measured blood glucose by Contour blood Glucose meter (Bayer) on day 7 post-infection.

